# Preference and switching in the Kill-the-Winner functional response: Diversity, size structure, and synergistic grazing in plankton models

**DOI:** 10.1101/848564

**Authors:** Kevin M. Archibald, Heidi M. Sosik, Michael G. Neubert

**Affiliations:** Biology Department, Woods Hole Oceanographic Institution, Woods Hole, MA, USA; Department of Earth, Atmospheric, and Planetary Sciences, Massachusetts Institute of Technology, Cambridge, MA, USA

## Abstract

Grazing by zooplankton can maintain diversity in phytoplankton communities by allowing coexistence between competitors in situations that would otherwise lead to competitive exclusion. In mathematical models, grazing is represented by a functional response that describes the consumption rate by an individual zooplankter as a function of phytoplankton concentration. Since its initial description, the Kill-the-Winner functional response has been increasingly adopted for large-scale biogeochemical modeling. Here, we analyze how two properties of the Kill-the-Winner functional response—preference and switching—interact to promote coexistence and increase diversity in two simple models: a diamond-shaped nutrient-phytoplankton-zooplankton model and a size-structured phytoplankton community model. We found that, compared to preference, switching leads to coexistence and increased diversity over a much wider range of environmental conditions (nutrient supply and mixing rate). In the absence of switching, preference only allows for coexistence within the narrow range of environmental conditions where the preference is precisely balanced against the competitive difference between phytoplankton types. We also explored a counterintuitive aspect of the Kill-the-Winner functional response that we have termed “synergistic grazing”. Synergistic grazing occurs when the grazing rate on one phytoplankton type increases as the biomass of an alternative phytoplankton type increases. This unrealistic effect is most evident when switching is strong and when zooplankton have a preference for the weaker competitor.

## Introduction

Phytoplankton communities commonly consist of many species living together in an apparently homogeneous environment and competing for a small number of limiting resources. This coexistence perplexed ecologists in the middle of the twentieth century because the current understanding of ecology led them to believe that the strongest competitor should exclude all others [1]. Hutchinson [2] termed this disparity between observations and ecological theory the “paradox of the plankton.” Beginning in the 1960s and 70s, a number of ecological mechanisms were identified that promote species coexistence and counteract the competitive exclusion principle. These include fluctuating environmental conditions that prevent the community from reaching equilibrium [3], multiple limiting resources [4], and, the object of our attention, the top-down effects of grazing [5]. In particular, it has been shown that grazers switching between multiple prey species can mediate coexistence between competitors and increase diversity in food webs [6–8].

Mathematical models have proven to be a useful tool in studying how grazing can modulate competitive interactions and mediate coexistence [9–12]. In models, grazing is determined by the *functional response*, the relationship between the amount of a resource and the rate that that resource is consumed by a single grazer. Holling [13] described three types of functional responses that have become standard in ecological models: linear, concave, and sigmoid (commonly referred to as Types I, II, and III). These classic functional responses can be derived from first principles based on the attack rate and handling time of the grazer for a given resource. However, functional responses that include multiple resources are more complicated because the relationship between consumption rate and resource density may be different for different resources. These relationships may be further complicated by the fact that the consumption rate of one resource may be affected by the abundance of other available resources [14, 15]. Gentleman et al. [16] provide a review of functional responses that have been used in the literature to describe zooplankton grazing on multiple phytoplankton types.

The relationship between zooplankton grazers and different phytoplankton types in the environment can be complex. When considering different mechanisms by which zooplankton may promote coexistence between competing phytoplankton, it is helpful to be precise about terminology for specific grazer characteristics. A zooplankter is said to exhibit *preference* for a phytoplankton type when the proportion of that type in its diet exceeds the proportion of that type in the environment [12, 17, 18]. A zooplankter is said to exhibit *switching* when the proportion of a phytoplankton type in its diet changes from less than expected to greater than expected as the proportion of that phytoplankton type in the environment increases [18]. Functional responses that incorporate preference and switching have been included in models to test whether these behaviors promote coexistence between competing phytoplankton species or size classes in circumstances that would otherwise lead to competitive exclusion [16, 19]. Global biogeochemical models with these kinds of functional responses show increased phytoplankton diversity compared to those that do not include these behaviors [8, 20]. Preference and switching are commonly included simultaneously in these models, however, and the individual impacts of each characteristic are rarely considered.

Functional responses that describe zooplankton grazing on multiple phytoplankton types are usually phenomenological and their parameters often cannot be measured directly. Some of these functional responses have characteristics that are biologically unrealistic [16]. One such characteristic is *antagonistic feeding*, in which a zooplankter has a higher total consumption rate when feeding on a single phytoplankton type than it would when the same total phytoplankton biomass is divided among many different types [21]. To avoid antagonistic feeding, Vallina et al. [20] developed an alternative formulation, the Kill-the-Winner (KTW) functional response. Since its publication, the KTW functional response has become a popular way to include switching behavior in a variety of models [22–32].

Vallina et al. [20] used one case of the KTW functional response to study how grazer switching promotes diversity on a global scale in a size-structured phytoplankton community model. Their model included the transport of plankton via a global circulation model. Here, we eliminate spatial dispersal in order to isolate and focus on biological interactions on the scale of competition between individual phytoplankton types. We have broken down our analysis to consider the capacity of zooplankton preference and switching as independent mechanisms for mediating coexistence between competing phytoplankton. We also identify a specific characteristic of the KTW functional response, which we refer to as “synergistic grazing”, that can generate counterintuitive results under certain conditions.

## Model Construction and Analysis

### Preference and switching in the KTW functional response

Whenever multiple phytoplankton types are available for consumption, a zooplankter’s consumption rate on a particular type may depend on its preference for that type. Zooplankton preference may arise from differential searching rates or rejection of less desirable phytoplankton types [17]. By definition, preference is fixed and independent of the distribution of phytoplankton types in the environment [18]. A zooplankter’s consumption rate on a given type may also depend upon the abundance of other available types in the environment via switching. Switching represents some behavioral change in the zooplankton that occurs in response to a variable phytoplankton community. Such responses include changing feeding strategies or learning how to capture or handle a particular phytoplankton type more efficiently [18, 20].

The KTW functional response includes both preference and switching. It defines the per capita grazing rate on phytoplankton type *P*_*i*_ as

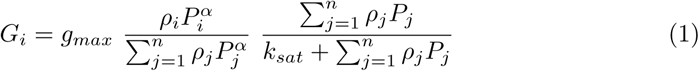

[20]. *G*_*i*_ has two components. One component, 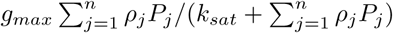, represents the total grazing rate as it depends upon the total preference-weighted biomass of the *n* phytoplankton types. It takes the form of a typical Holling Type II functional response with a maximum consumption rate *g*_*max*_ and half-saturation constant *k*_*sat*_. The preference for phytoplankton type *i* is *ρ*_*i*_. If the zooplankter has no preference for any phytoplankton type then all the values of *ρ*_*i*_ are equal. If the zooplankter does have a preference for a phytoplankton type, then the values of the *ρ*_*i*_ will not all be equal; a larger value of *ρ*_*i*_ indicates a stronger preference for type *i*.

The other component, 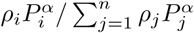, specifies the portion of the total grazing rate that is applied to the *ith* phytoplankton type. The value of *α* controls the switching behavior. If the zooplankter does not exhibit switching, then *α* = 1. When *α* > 1, the zooplankter exhibits some degree of switching with larger values of *α* corresponding to stronger switching behaviors. For very large values of *α*, the grazing pressure is concentrated entirely on the most abundant phytoplankton type. In prior work, the KTW functional response has only been examined for *α* = 2 [20].

If the zooplankter exhibits neither preference nor any switching behavior, then the proportion of a phytoplankton type in the zooplankter’s diet will be equal to that type’s proportion in the environment (Fig 1, solid black line). If the zooplankter exhibits preference for a phytoplankton type, but no switching, then the proportion of a phytoplankton type in the zooplankter’s diet will be higher than its proportion in the environment (Fig 1, solid blue line). If the zooplankter exhibits switching, then the proportion of a phytoplankton type in the zooplankter’s diet will change from less than expected to greater than expected (relative to the case where the zooplankter has no preference or switching) as the proportion of that type in the environment increases (Fig 1, dashed lines). Switching to the more common phytoplankton type occurs at a lower proportional abundance if the zooplankter has a preference for that type, and at a higher proportional abundance if it has a preference for other types.

**Fig 1.**
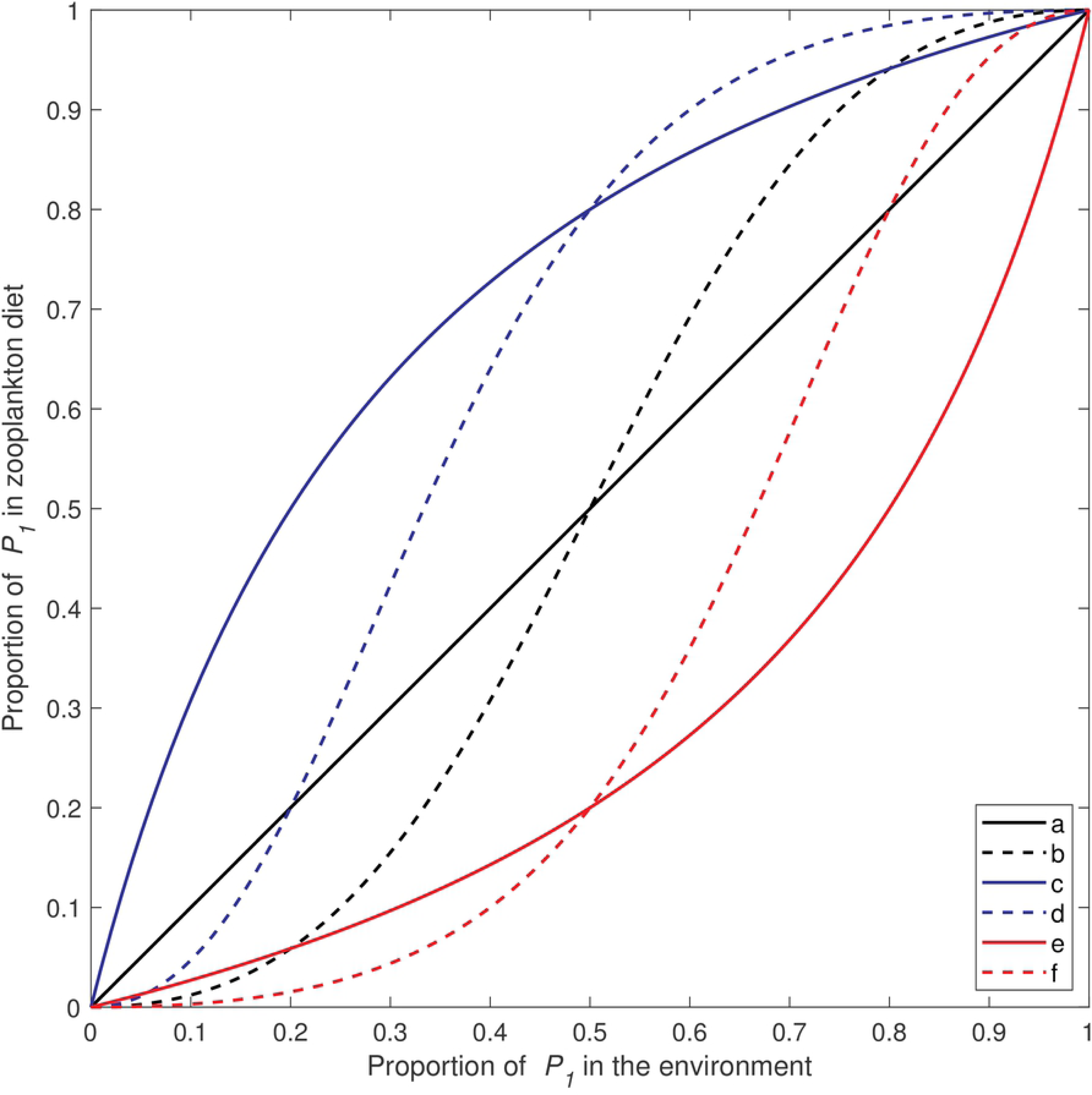
Characteristics of the KTW functional response using different combinations of preference and switching. The proportion of *P*_1_ in a zooplankter’s diet as a function of the proportion of *P*_1_ in the environment based on Eq 1 and assuming two phytoplankton types. The different curves show different combinations of preference and switching: (a) neither preference nor switching (*ρ*_1_ = *ρ*_2_, *α* = 1), (b) switching without preference (*ρ*_1_ = *ρ*_2_, *α* > 1), (c) preference for *P*_1_ without switching (*ρ*_1_ > *ρ*_2_, *α* = 1), (d) preference for *P*_1_ with switching (*ρ*_1_ > *ρ*_2_, *α* > 1), (e) preference for *P*_2_ without switching (*ρ*_1_ *< ρ*_2_, *α* = 1), (f) preference for *P*_2_ with switching (*ρ*_1_ *< ρ*_2_, *α* > 1).

### The diamond-shaped food web model

Switching and preference can have dramatic effects on the dynamics of plankton food web models. As a relatively simple example, consider the following diamond-shaped food web with one nutrient resource (*N*), two competing phytoplankton types (*P*_1_ and *P*_2_), and a single zooplankton species (*Z*) that feeds upon both phytoplankton. The model assumes a constant and well-mixed environment, such as the mixed layer in a pelagic water column. Nutrients are supplied to the mixed layer through vertical mixing with the deep ocean. This influx water has a nutrient concentration *N*_0_ and is supplied at a rate *D*. Nutrients and phytoplankton are mixed out of the water column at the same rate *D*. We assume that zooplankton are relatively strong swimmers and are not mixed out. The maximum growth rates of *P*_1_ and *P*_2_ are given by *µ*_1_ and *µ*_2_. Nutrient uptake by the phytoplankton follows Monod kinetics with half-saturation constants *k*_1_ and *k*_2_. The phytoplankton non-grazing mortality rate is given by *m*_*p*_ and the mortality of the zooplankton is given by *m*_*z*_. *γ* is the growth efficiency of the zooplankton. The dynamics of this food web are given by

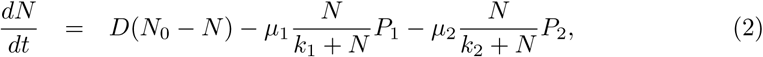

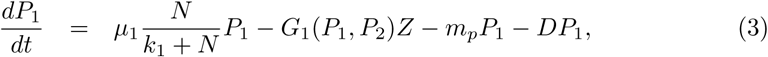

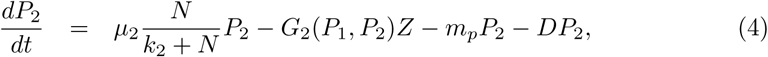

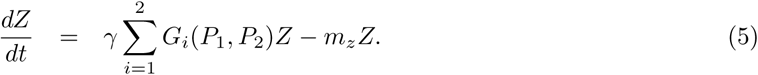

#### Coexistence dynamics in the diamond-shaped food web model

In the absence of any grazing (i.e., when *Z* = 0), the phytoplankton type that is the better competitor for nutrients will eventually exclude the other phytoplankton type in model (2)-(5) [33]. We define the value

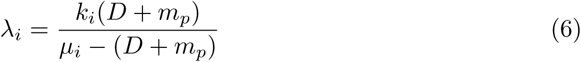

as a metric for the competitive ability of a phytoplankton type. If *λ*_*i*_ *< λ*_*j*_, then *P*_*i*_ is a stronger competitor for nutrients than *P*_*j*_ and *P*_*i*_ will drive *P*_*j*_ to extinction [33]. We chose *µ*_1_ > *µ*_2_ and *k*_1_ *< k*_2_ to satisfy the criteria that *P*_1_ is the stronger competitor (Table 1). Zooplankton preference and switching may allow for coexistence between *P*_1_ and *P*_2_ in situations where, in the absence of the zooplankton, *P*_1_ would drive *P*_2_ to extinction. Consider the scenario where *P*_2_ = 0; the equilibrium point of the linear food chain composed of *N, P*_1_, and *Z* is given by

**Table 1.** Parameter values used to simulate the diamond-shaped food web model (2)-(5)

| Parameter | Description | Value | Units |
| --- | --- | --- | --- |
| $N_0$ | influx water nutrient concentration | (1, 15) | $mmol\ N\ m^{-3}$ |
| $D$ | mixing rate | $(10^{-4}, 10^{-1})$ | $d^{-1}$ |
| $\mu_1$ | maximum growth rate of $P_1$ | 1.0 | $d^{-1}$ |
| $\mu_2$ | maximum growth rate of $P_2$ | 0.8 | $d^{-1}$ |
| $k_1$ | $P_1$ nutrient uptake half-saturation constant | 0.1 | $mmol\ N\ m^{-3}$ |
| $k_2$ | $P_2$ nutrient uptake half-saturation constant | 0.15 | $mmol\ N\ m^{-3}$ |
| $m_p$ | non-grazing phytoplankton mortality rate | 0.01 | $d^{-1}$ |
| $g_{max}$ | maximum grazing rate | 2.0 | $d^{-1}$ |
| $k_{sat}$ | grazing half-saturation constant | 10 | $mmol\ N\ m^{-3}$ |
| $\gamma$ | zooplankton growth efficiency | 0.7 | |
| $m_z$ | zooplankton mortality rate | 0.1 | $d^{-1}$ |

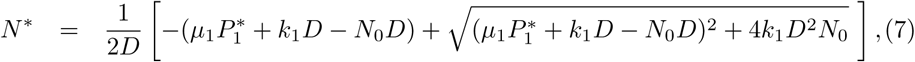

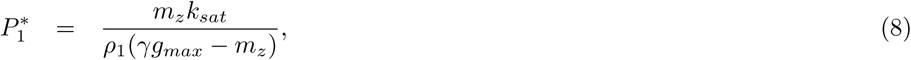

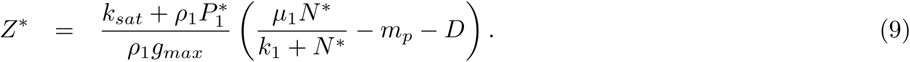

Near this equilibrium point, the dynamics of *P*_2_ are given by the linearization of model (2)-(5). The population of type *P*_2_ will grow (or shrink) according to the invasion growth rate *r*—the per capita growth rate of *P*_2_ when *P*_2_ ≈ 0:

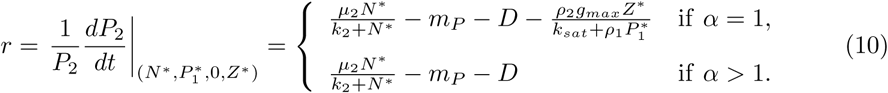

If *r* > 0, then *P*_2_ can successfully invade the system.

Note that the term 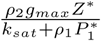 in Eq 10 is always positive, so the growth of a phytoplankton type when its density is near zero is strictly greater when the zooplankton grazer displays switching. Switching behavior, therefore, is a kind of density-dependent mortality that provides a refuge for phytoplankton at low densities. Eq 10 suggests that there exist some values of *ρ*_*i*_ and *α* such that *r* > 0 even when *λ*_2_ *< λ*_1_. Stated differently, a weaker phytoplankton competitor should be able to invade the equilibrium of a stronger phytoplankton competitor under certain grazing conditions. We explored this hypothesis further by simulating the diamond-shaped food web model in each of six test cases which cover different combinations of preference and switching.

Case 1: control scenario, no preference (*ρ*_1_ = *ρ*_2_ = 1) and no switching (*α* = 1)

Case 2: preference for *P*_1_ (*ρ*_1_ = 1, *ρ*_2_ = 0.5) without switching (*α* = 1)

Case 3: preference for *P*_2_ (*ρ*_1_ = 0.5, *ρ*_2_ = 1) without switching (*α* = 1)

Case 4: no preference (*ρ*_1_ = *ρ*_2_ = 1) with switching (*α* = 2)

Case 5: preference for *P*_1_ (*ρ*_1_ = 1, *ρ*_2_ = 0.5) with switching (*α* = 2)

Case 6: preference for *P*_2_ (*ρ*_1_ = 0.5, *ρ*_2_ = 1) with switching (*α* = 2)

For each case, we simulated model (2)-(5) over a range of input nutrient concentrations (*N*_0_) and mixing rates (*D*) using random initial conditions drawn from uniform distributions. We observed four possible asymptotic outcomes: (1) *P*_1_ drives *P*_2_ to extinction, (2) *P*_2_ drives *P*_1_ to extinction, (3) *P*_1_ and *P*_2_ exist in a stable equilibrium, or (4) *P*_1_ and *P*_2_ exist in an unstable equilibrium characterized by oscillations. For these simulations, *P*_1_ was the stronger competitor and *P*_2_ was the weaker competitor.

These simulations suggest that coexistence is possible in the diamond-shaped food web model if (1) the zooplankton have a preference for the stronger competitor or (2) the zooplankton exhibit switching (Fig 2). However, the range of environmental conditions (i.e., nutrient inputs and mixing rates) that allows for coexistence in Cases 2 and 3, where zooplankton exhibit preference only, is quite small, indicating that coexistence mediated by preference alone is rare and represents a delicate balance between zooplankton preference and the growth rates and nutrient affinities of the competing phytoplankton types. In contrast, for Cases 4-6, where zooplankton exhibit switching, coexistence occurs over a wider range of environmental conditions, indicating that switching is a robust mechanism for promoting coexistence. Finally, there was only one case in which the weaker phytoplankton type drove the stronger phytoplankton type to extinction (Fig 2). When zooplankton have a preference for the stronger phytoplankton and display no switching (Case 2), *P*_2_ > 0 and *P*_1_ = 0 at equilibrium. The simulation results were consistent with our predictions based on Eq 10. The *r* = 0 line where the invasion growth rate for the weaker competitor changes from negative to positive corresponds to the boundary between the region of the parameter space where the model is asymptotically stable with *P*_2_ = 0 and the region where stable coexistence between *P*_1_ and *P*_2_ occurs (Fig 2).

**Fig 2.**
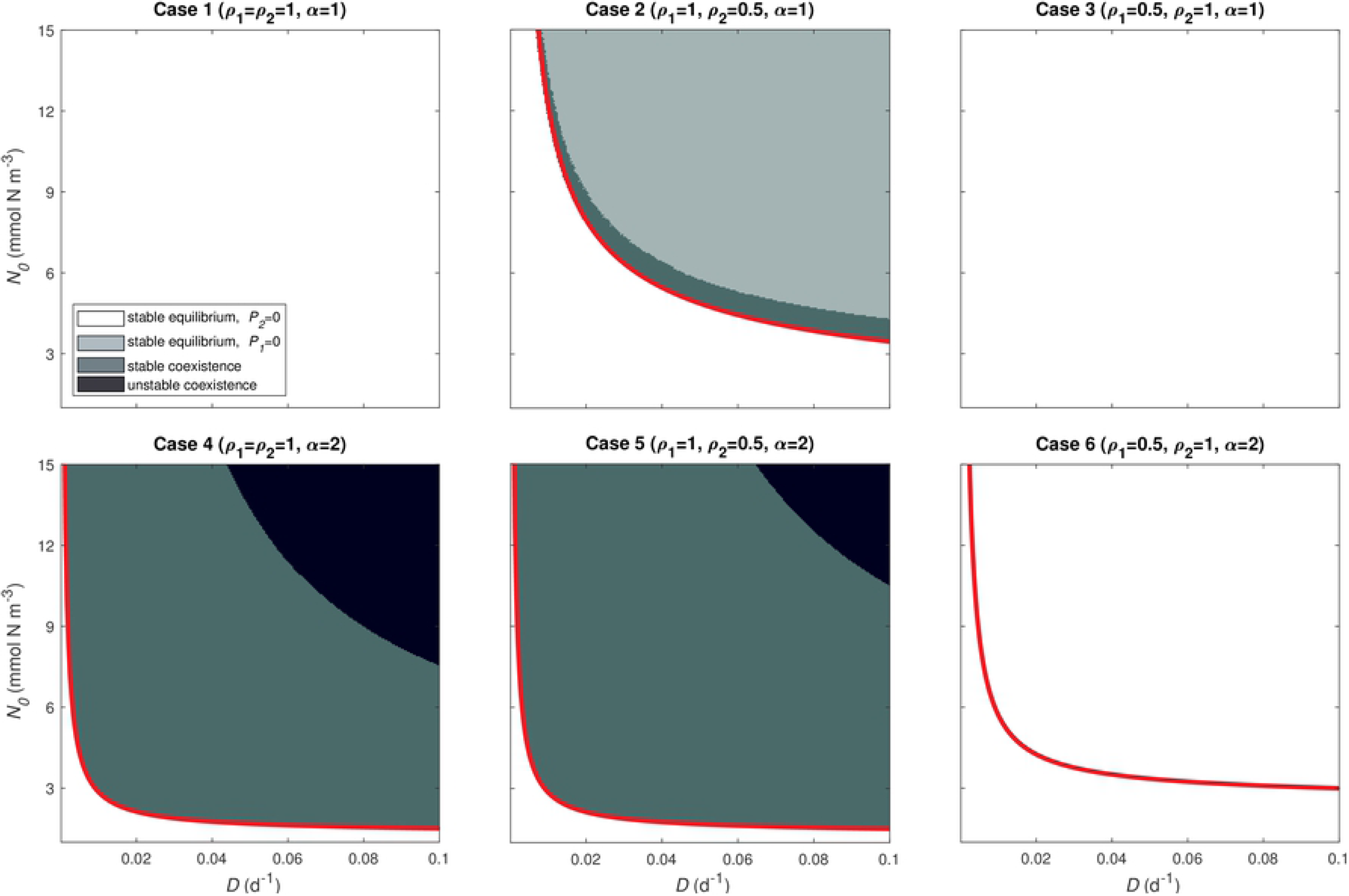
Asymptotic dynamics of the diamond-shaped food web model. The six subplots show the asymptotic behavior of model (2)-(5) as a function of *N*_0_ and *D* for Cases 1-6 listed above. For each parameter combination we chose initial conditions at random from uniform distributions (*N* (0) ∼ *U* [0, 15], *P*_1_(0) ∼ *U* [0, 2], *P*_2_(0) ∼ *U* [0, 2], *Z*(0) ∼ *U* [0, 10]), and simulated the model from *t* = 0 to *t* = 10, 000 days. Colors indicate the asymptotic result. The red line for Cases 2 and 4-6 shows the root of Eq 10 as a function of *N*_0_ and *D*. Below the line, *r <* 0 and above the line *r* > 0.

#### Synergistic grazing in the diamond-shaped food web model

The KTW functional response has one characteristic that bears particular attention. The grazing rate on one phytoplankton type can increase as the density of an alternative phytoplankton type increases (Fig 3). Here, we coin the term *synergistic grazing* to describe this phenomenon. Mathematically, synergistic grazing occurs when 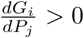 for *i* ≠ *j*. Synergistic grazing is not unique to the KTW functional response [16].

**Fig 3.**
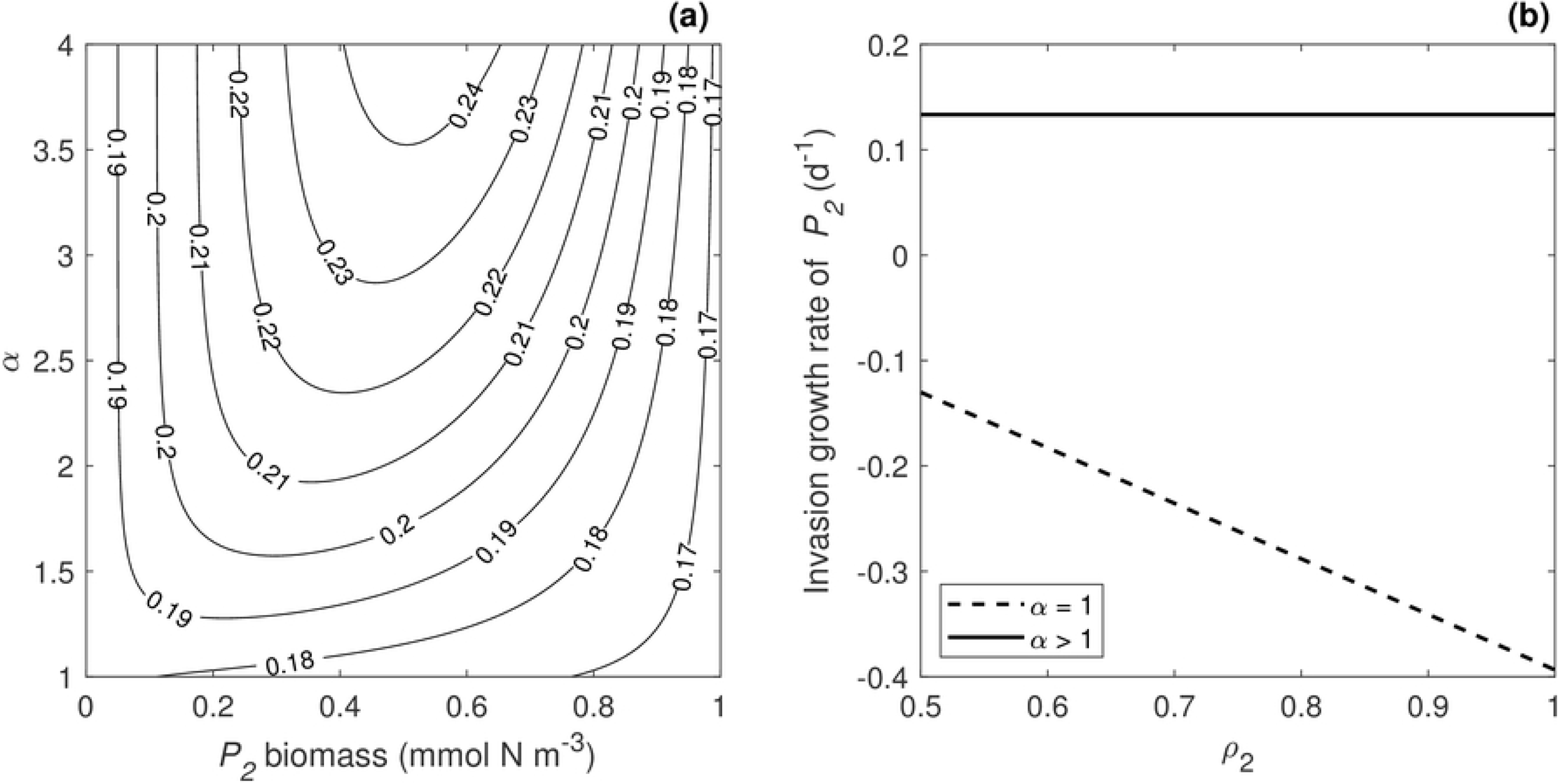
Synergistic grazing in the diamond-shaped food web model. (a) Zooplankton grazing rate on *P*_1_ as a function of *α* and *P*_2_ and (b) the invasion growth rate of *P*_2_ as function of *ρ*_2_ for *α* = 1 (no switching) and *α* > 1 (switching). In panel (b), *ρ*_1_ = 0.5.

In the KTW functional response, zooplankton total ingestion rate depends on the total available phytoplankton biomass independent of the proportions of different phytoplankton types (Eq 1). The ingestion rate of individual phytoplankton types, or the distribution of the total grazing pressure, depends on the proportion of a phytoplankton type in the environment and on the zooplankton preference.

Consider the scenario in which we begin with only one phytoplankton type (*P*_1_) and we calculate the grazing rate on *P*_1_ as we introduce a secondary phytoplankton type (*P*_2_). The total grazing rate must increase because the total phytoplankton biomass is increasing. However, if switching is strong (*α* >> 1), then the distribution of grazing pressure will be heavily skewed towards *P*_1_ since *P*_2_ exists at low densities in the environment. In this case, the grazing rate on *P*_1_ will increase as *P*_2_ is introduced even though the density of *P*_1_ is being held constant. Perhaps counterintuitively, synergistic grazing has a stronger effect when the predator has a strong preference for the less abundant phytoplankton type because the value of *ρ*_*i*_ acts as a modifier of the “effective biomass” in the system. Therefore, if *ρ*_2_ is large, then more effective biomass is added to the system when *P*_2_ is introduced and the grazing rate changes faster than it would if the zooplankton had a lower preference for *P*_2_.

Synergistic grazing alters the dynamics of the diamond-shaped food web model in interesting ways. It has a larger effect when switching behavior is stronger (Fig 3) and in some cases can result in coexistence criteria that are counterintuitive. For example, in a model with no switching, the invasion growth rate of *P*_2_ will decrease as the zooplankton preference for *P*_2_ increases (Fig 3). However, when switching is included, the invasion growth rate is independent of *ρ*_2_, meaning that *P*_2_ will be able to invade the system no matter how large the grazer preference for *P*_2_ becomes. This occurs because switching creates a grazing refuge for phytoplankton at low densities. Grazer preference has no effect when a phytoplankton type is rare because all of the grazing is focused on the more common phytoplankton type.

### The size-structured phytoplankton model

Phytoplankton communities rarely exist as only two types, but instead are commonly composed of many different types coexisting simultaneously. Size is a particularly important trait for structuring these communities since cell size is related to many important characteristics of phytoplankton, including growth rates and nutrient affinities [45]. To explore the effects of zooplankton preference and switching on phytoplankton communities structured by cell size, we extended the model above to an NPZ model that includes an arbitrary number of phytoplankton size classes. This new model maintains a single zooplankton species that distributes grazing pressure across all phytoplankton size classes according to the KTW functional response. The size-structured model is written as

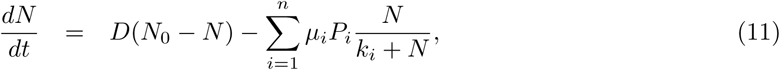

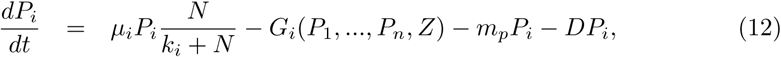

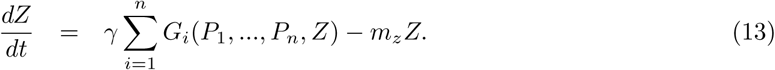

The parameters *µ*_*i*_ and *k*_*i*_ are the growth rate and nutrient affinity for each phytoplankton size class. We chose to scale these parameters allometrically (see Table 2) such that the smallest phytoplankton size classes have the largest *µ* and the smallest *k* following previous size-structured phytoplankton models [34, 35]. As size increases, competitive ability decreases. Other parameters used to simulate the model are similar to those selected for the diamond-shaped model (Table 2).

**Table 2.** Parameter values used to simulate the size-structured phytoplankton community model.

| Parameter | Description | Value | Units |
| --- | --- | --- | --- |
| $N_0$ | influx water nutrient concentration | (1, 15) | $mmol\ N\ m^{-3}$ |
| $D$ | mixing rate | $(10^{-4}, 10^{-1})$ | $d^{-1}$ |
| $n$ | number of phytoplankton size classes | 10 | |
| $x$ | cell volume | $\log_{10} x = [-1 + \frac{3}{n-1}(i-1)], i = 1 : n$ | $\mu m^3$ |
| $\mu$ | maximum phytoplankton growth rate | $0.5x^{-0.24}$ | $d^{-1}$ |
| $k$ | nutrient uptake half-saturation constant | $0.14x^{0.33}$ | $mmol\ N\ m^{-3}$ |
| $m_p$ | non-grazing phytoplankton mortality rate | 0.01 | $d^{-1}$ |
| $g_{max}$ | maximum grazing rate | 2.0 | $d^{-1}$ |
| $k_{sat}$ | grazing half-saturation constant | 10 | $mmol\ N\ m^{-3}$ |
| $\gamma$ | zooplankton growth efficiency | 0.7 | |
| $m_z$ | zooplankton mortality rate | 0.1 | $d^{-1}$ |
| $\alpha$ | zooplankton switching strength | (1,3) | |

Our analysis of the diamond-shaped food web model indicated that coexistence mediated by preference alone was possible for only a small set of environmental conditions. During preliminary analysis of the size-structured model, we found a similar result — namely, that preference alone would result in coexistence only if grazer preference was carefully balanced against the competitive imbalance between phytoplankton size classes. Additionally, this balance was significantly more difficult to achieve in the size-structured model due to the increased number of phytoplankton types compared to the diamond-shaped food web model. For this reason, we have chosen to focus on the scenarios that include switching, which allows coexistence over a much broader range of parameter values and environmental conditions compared to preference alone. We defined three new cases that all included zooplankton switching (*α* = 2).

Case 7: no preference for any size class

Case 8: zooplankton preference for smaller phytoplankton size classes

Case 9: zooplankton preference for larger phytoplankton size classes

The distribution of grazer preference between size classes is more complex in model (11)-(13). For Case 7 (no preference), by definition, *ρ*_*i*_ is equal across all size classes. For Case 8 (preference for smaller size classes), we set *ρ*_*i*_ to decrease linearly across size classes. For Case 9 (preference for larger size classes), we set *ρ*_*i*_ to increase linearly across size classes. Taking *ρ*_1_ and *ρ*_*n*_ as the preference for the largest and smallest size classes, then *ρ*_*i*_ for each size class is

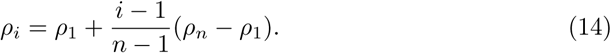

Previous analysis of the KTW functional response has assumed a constant *α* = 2 [20]. With that same assumption, we find that coexistence between at least two phytoplankton size classes is possible in Cases 7 and 8, as long as the influx water nutrient concentration (*N*_0_) is large enough (Fig 4). In Case 9, the smallest phytoplankton size class dominated and excluded all other size classes. In all of these simulations, the smallest phytoplankton size class is always the only one to exist by itself at low nutrient input rates and larger size classes are added in order of increasing size as *N*_0_ increases (Fig 4).

**Fig 4.**
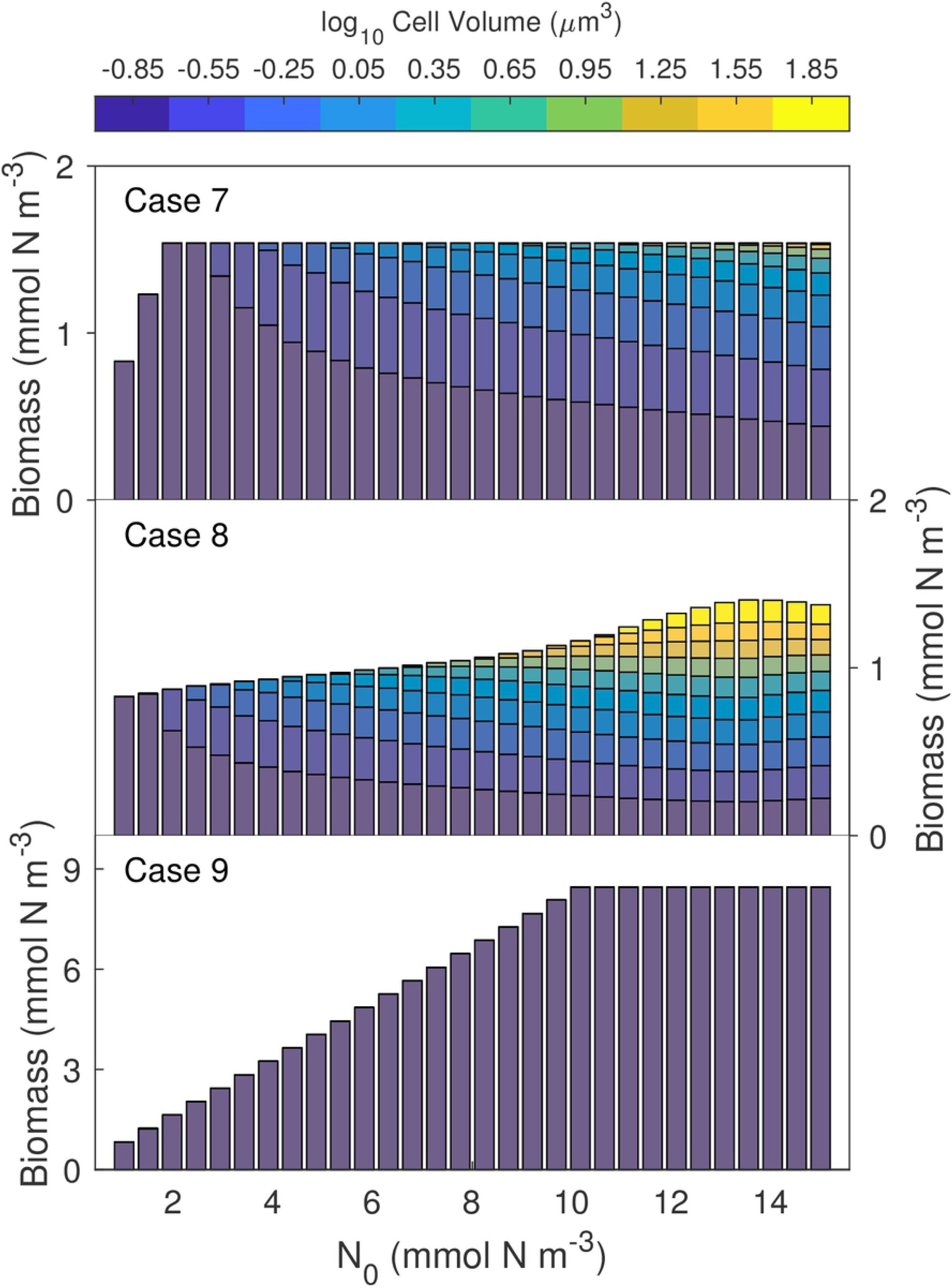
Simulations of the size-structured phytoplankton community model. We simulated model (11)-(13) with ten phytoplankton size classes for 30 values of *N*_0_ ranging from 1-15 *mmolNm*^−3^. We chose initial conditions at random from uniform distributions (*N* (0) ∼ *U* [0, 15], *P*_*i*_(0) ∼ *U* [0, 2], *Z*(0) ∼ *U* [0, 10]), and simulated the model from *t* = 0 to *t* = 10, 000 days. Each bar represents the steady-state phytoplankton community for a given value of *N*_0_. The height of the bar gives the total phytoplankton biomass and the bar is color coded by the cell volume of a given size class. Case 7 includes switching without preference (*ρ*_1_ = *ρ*_10_ = 1, *α* = 2), Case 8 includes switching with preference for smaller size classes (*ρ*_1_ = 1, *ρ*_10_ = 0.5, *α* = 2), and Case 9 includes switching with preference for larger size classes (*ρ*_1_ = 0.5, *ρ*_10_ = 1, *α* = 2).

We further explored how variable switching strength affected diversity by simulating the size-structured model for *α* = 1 to 3. In these simulations, we also considered a range of relative preference for smaller size classes (*ρ*_1_ − *ρ*_*n*_) where relative preference equals zero corresponds to Case 7 and values greater than zero correspond to Case 8 (Fig 5). Phytoplankton diversity, as represented by the number of coexisting size classes at steady state, is higher under stronger zooplankton switching and increased relative preference for smaller phytoplankton size classes. For Case 9, where zooplankton have preference for larger size classes, we found a counterintuitive relationship between preference and its impacts on phytoplankton size structure; we treat this in more detail in the next section.

**Fig 5.**
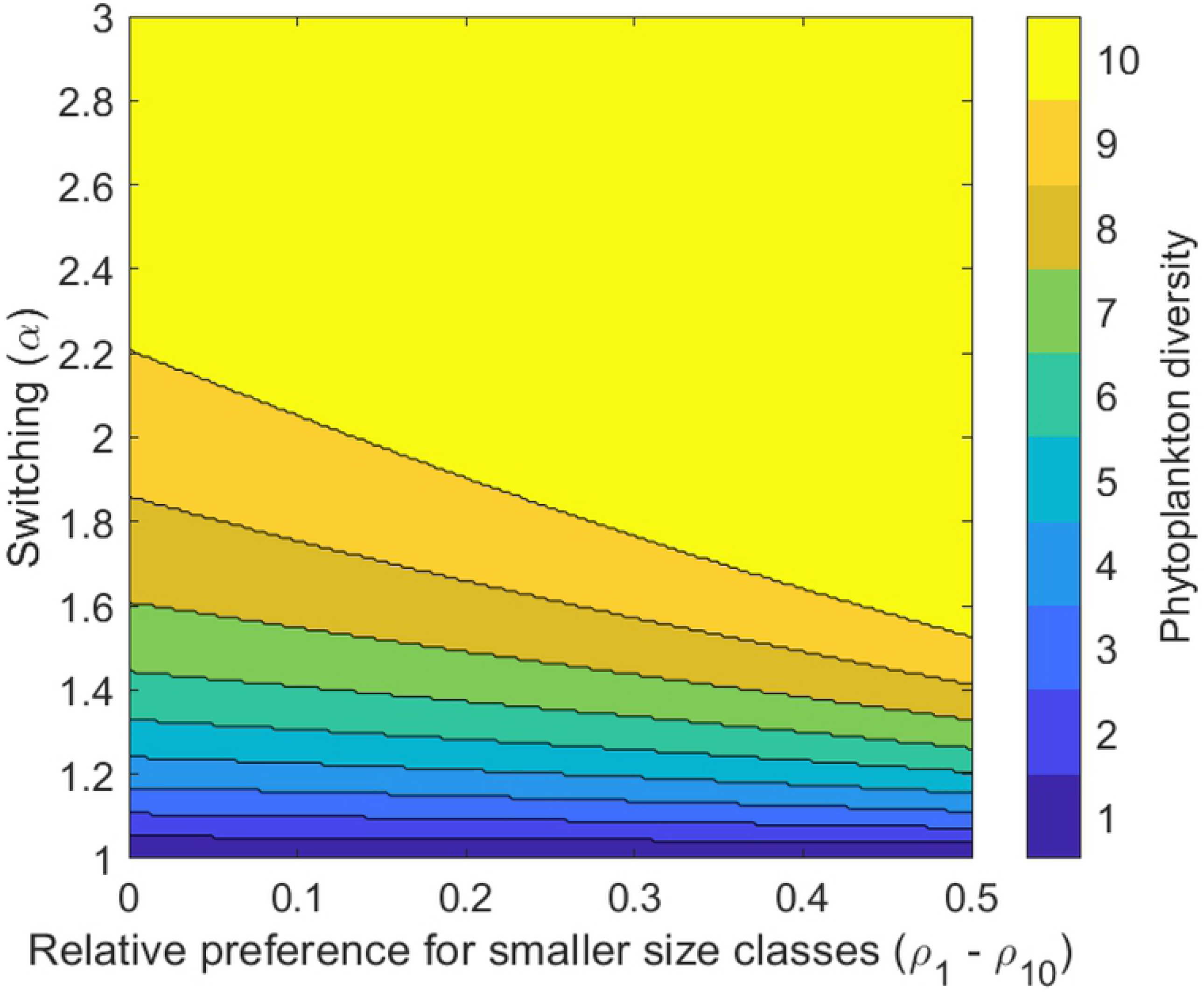
Zooplankton switching and preference for smaller phytoplankton size classes increase phytoplankton diversity. Phytoplankton diversity in model (11)-(13) as a function of the strength of zooplankton switching (*α*) and the difference between zooplankton preference for the smallest and largest size classes (*ρ*_1_ − *ρ*_10_). A larger difference indicates a stronger relative preference for smaller phytoplankton size classes.

#### Synergistic grazing in the size-structured model

The counterintuitive effects of synergistic grazing emerge when zooplankton prefer larger phytoplankton size classes (Case 9). To emphasize this effect, we simulated the size structured model using very strong switching, because the effects of synergistic grazing are more evident when *α* is larger. Intuitively, we expect that increased zooplankton preference for large size classes will make it harder for these size classes to persist, since larger cells are weaker competitors for nutrients to begin with. However, because of the effects of synergistic grazing in the KTW functional response, we actually found that larger size classes benefited from increased grazer preference when switching was very strong.

We ran a series of simulations of the size-structured model using *α* = 10 and different values of relative preference for larger size classes (*ρ*_*n*_ − *ρ*_1_) where relative preference equals zero corresponds to Case 7 and positive values correspond to Case 9 (Fig 6). Contrary to our expectation, an increased relative preference for larger size classes resulted in higher biomass within the largest phytoplankton size class and lower biomass in the smallest size class. The response in the intermediate size classes was complex, resulting in some cases in non-monotonic changes in the size class biomass as the relative preference was adjusted.

**Fig 6.**
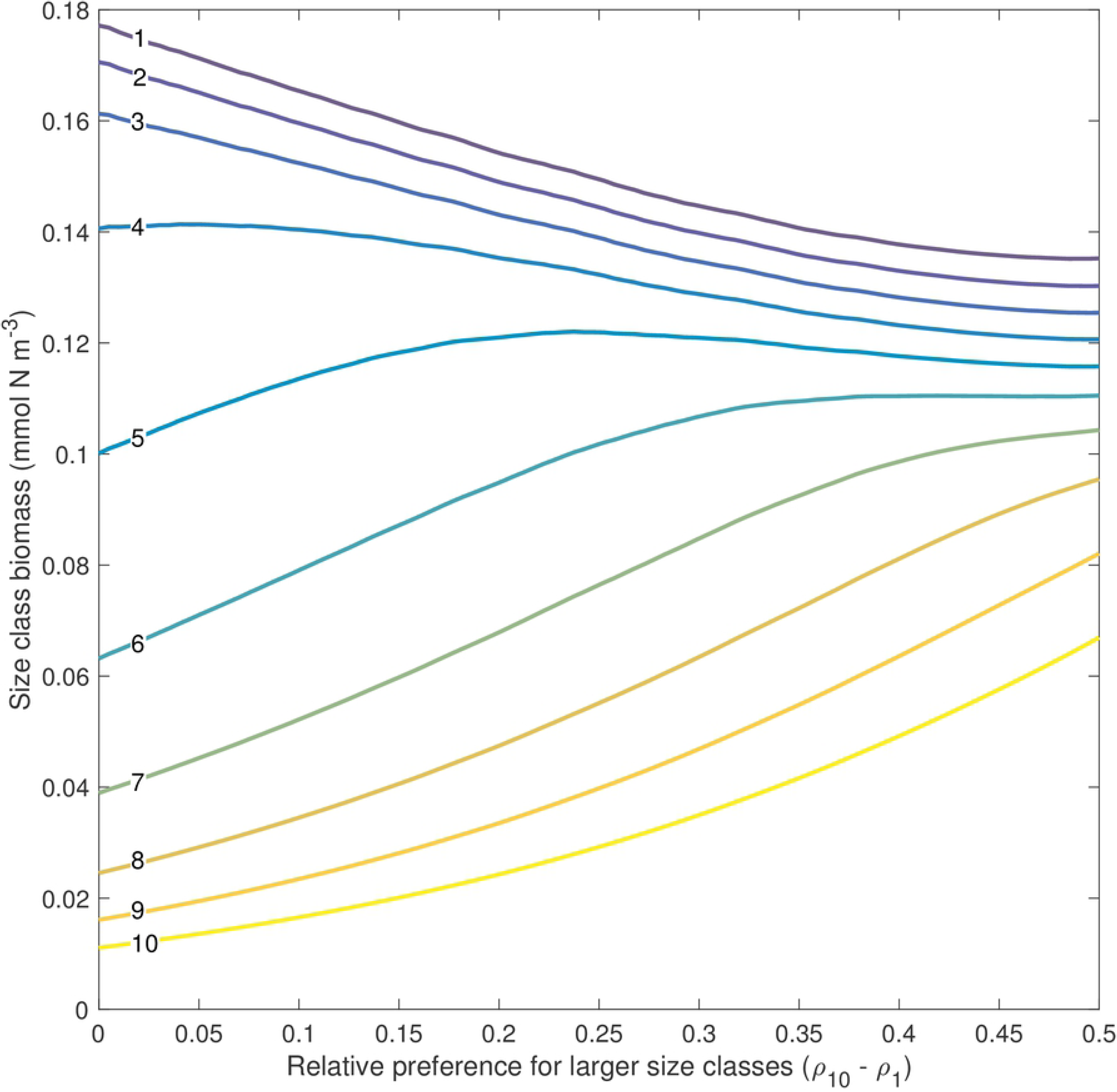
Synergistic grazing in the size-structured phytoplankton community model. Mean biomass concentration in each of the ten phytoplankton size classes over the steady-state limit cycle as a function of the difference between zooplankton preference for the largest and smallest size classes (*ρ*_10_ − *ρ*_1_). A larger difference indicates a stronger relative preference for larger phytoplankton size classes. Each line is labeled, with one referring to the smallest size class and ten referring to the largest size class. For these simulations, *α* = 10.

Strong switching results in grazing pressure that is heavily skewed towards the smaller size classes since they compose the majority of the phytoplankton biomass. This creates a scenario in which the grazing pressure on large size classes is low (due to their very low densities), even when there is strong preference for large size classes. Instead, strong preference for larger size classes increases the total preference-weighted biomass in the system, resulting in a higher overall grazing rate. A higher grazing rate actually favors the larger size classes since strong switching ensures that most of the grazing is focused on the smaller, more abundant phytoplankton types.

## Discussion

Predator switching behavior is frequently observed in laboratory and field studies [14, 36–39]; modeling work has shown that it stabilizes dynamics and increases diversity [8, 19, 20, 40]. Gentleman et al. [16] provide a thorough review of the functional responses commonly used in biogeochemical models, many of which include grazer preference and switching. Here, we analyzed the KTW functional response to evaluate the mechanisms by which preference and switching allow coexistence under circumstances that would otherwise lead to competitive exclusion. We have extended the work of Vallina et al. [20] by examining how the functional response behaves on the scale of competition between individual phytoplankton types. We have also identified and discussed an important characteristic of the functional response, which we have termed “synergistic grazing.” Synergistic grazing can result in biologically counterintuitive dynamics, particularly when strong switching is combined with preference for weaker competitors.

We found that the KTW functional response allowed for grazer-mediated coexistence through both grazer preference and switching as independent mechanisms. Switching was generally the more powerful mechanism for generating coexistence, allowing for coexistence between competing phytoplankton types across a broad set of environmental conditions. In contrast, grazer preference in the absence of switching must be carefully balanced with the competitive ability of the phytoplankton types to allow for coexistence. When combining both zooplankton preference and switching in the KTW parameterization, caution is warranted as the interaction can produce some unexpected behaviors. Synergistic grazing is an example of a potentially problematic characteristic that emerges as a result of the interaction between zooplankton preference and switching. The effects of synergistic grazing are most evident when switching is strong and when zooplankton have a preference for the weaker competitor. It is worth noting that synergistic grazing arises directly from the fact that total grazing rate and the distribution of the total grazing rate onto individual phytoplankton types are calculated as independent terms. This functional form was chosen specifically to avoid the problem of antagonistic grazing [20], another example of a problematic characteristic of grazing functional responses. Therefore, it appears that there is a trade-off between functional responses that display antagonistic grazing and the KTW functional response, which does not display antagonistic grazing, but does include synergistic grazing. To date, we are not aware of any functional response that includes a representation of switching behavior and displays neither antagonistic grazing nor synergistic grazing [16].

Conceptually, the switching functional response used in this model could represent multiple ecological mechanisms that all have the same emergent behavior: frequency-dependent differential grazing on multiple available phytoplankton types. The classical view of switching is that it arises from zooplankton behavior such as changes in feeding strategy or time budget optimization in response to a variable phytoplankton community [41–43]. Alternatively, the zooplankton represented in this model could be viewed as a generalized grazer community rather than a single species. Under this scenario, the switching “behaviors” represent internal changes in the composition of the zooplankton community as the composition of the phytoplankton community evolves through time. When one phytoplankton type becomes very abundant, specialized predators of that type also become more abundant and the integrated grazer community becomes more efficient at grazing that phytoplankton type. Conversely, when a phytoplankton type is rare, its specialized predators will also be rare and the proportion of total grazing pressure on this phytoplankton type will be small. From either perspective, the mathematical consequences are the same and so the parameterizations used here could be used to represent either ecological scenario.

Marine ecologists have long acknowledged the role that cell size plays in structuring phytoplankton communities [44]. Observations from a variety of diverse biogeochemical regimes have consistently shown an inverse relationship between cell size and phytoplankton abundance [45]. Furthermore, the fractional abundance of small cells in a phytoplankton community increases as total biomass decreases [45]. There has been some debate among ecologists as to exactly what environmental factors control phytoplankton community size structure, some claiming that nutrient supply alone is sufficient to explain the variability [46, 47], while others argue that temperature has an important effect [48]. Our analysis highlights another layer in this debate; grazing pressure, particularly when zooplankton display preference or switching, can play an important role in structuring phytoplankton communities. The inclusion of preference and switching behaviors increases phytoplankton diversity and restrains the dominance of smaller cells that, generally speaking, are stronger competitors for nutrients.

## Acknowledgments

Many thanks to Stephanie Dutkiewicz and Dennis McGillicuddy for discussions related to this work which have proven very helpful and increased the quality of this study. This work was supported by grants from NSF (OCE-1655686, HMS/MGN; OCE-1657803, HMS) and the Simons Foundation (561126, HMS).

